# Tamoxifen exacerbates morbidity and mortality in mice receiving medetomidine anaesthesia

**DOI:** 10.1101/2022.11.29.518236

**Authors:** Victoria S. Rashbrook, Laura Denti, Christiana Ruhrberg

## Abstract

Tamoxifen-induced CreER-LoxP recombination is often used to induce spatiotemporally controlled gene deletion in genetically modified mice. Prior work has shown that tamoxifen and tamoxifen-induced CreER activation can have off-target effects that should be controlled. However, it has not yet been reported whether tamoxifen administration, independently of CreER expression, interacts adversely with commonly used anaesthetic drugs such as medetomidine or its enantiomer dexmedetomidine in mice. Here we report a high incidence of urinary plug formation and morbidity in male mice on a mixed C57Bl6/J and 129/SvEv background when tamoxifen treatment was followed by ketamine-medetomidine anaesthesia. Medetomidine is therefore contra-indicated for male mice after tamoxifen treatment. As dexmedetomidine causes morbidity and mortality in male mice at higher rates than medetomidine even without tamoxifen treatment, our findings suggest that dexmedetomidine is not a suitable alternative for anaesthesia of male mice after tamoxifen treatment. We conclude that the choice of anaesthetic drug needs to be carefully evaluated in studies using male mice that have undergone tamoxifen treatment for inducing CreER-LoxP recombination.

## Introduction

The Cre-LoxP system is often used to induce gene deletion in mice, whereby the P1 bacteriophage Cre recombinase (hereafter referred to as Cre) is inserted together with a promoter into the mouse genome to excise the LoxP-flanked genetic material in cell types that express Cre ^1^. CreER is a modified version of Cre, in which Cre is fused to the oestrogen receptor (estrogen receptor, ER) to retain it in the cytoplasm, but CreER translocates to the nucleus after binding the tamoxifen metabolite 4-hydroxy tamoxifen (4-OHT) ^1^. Thus, treating mice with tamoxifen or 4-OHT enables temporally controlled gene deletion ^2, 3^.

Although CreER-mediated gene modification is now a widely used model to study gene function in the mouse, both tamoxifen and CreER induction have been reported to cause toxicity, with the extent of toxicity dependent on the tamoxifen dose, because tamoxifen itself can be toxic at high concentrations, but also because a higher tamoxifen dose induces more CreER nuclear translocation and thus more off-target effects ^4-7^. For example, the intraperitoneal injection of pregnant dams with tamoxifen at embryonic day (E) 9.75 caused a high incidence of limb abnormalities in E17 mouse embryos ^8^. Tamoxifen also causes intrauterine haemorrhage and increases the mortality of pregnant mice injected at E5.5 ^9^, but tamoxifen treatment of non-pregnant mice does not increase mortality ^9^.

CreER-modified transgenic mice may also be used in procedures that require anaesthesia after tamoxifen administration to induce gene deletion. Thus, the α2-adrenoreceptor antagonist medetomidine or its active enantiomer dexmedetomidine are commonly included in injectable anaesthetic regimes that are used in both veterinary medicine and research studies using live mice ^10^. However, it has not been reported whether tamoxifen exacerbates adverse effects of dexmedetomidine or medetomidine. Here we describe urinary plug formation after ketamine-medetomidine anaesthesia in tamoxifen-treated male mice. Although medetomidine causes morbidity and mortality in male mice at lower rates than at higher rates than dexmedetomidine ^11, 12^, our findings nevertheless suggest that medetomidine, even though preferable over dexmedetomidine, is contra-indicated for male mice after tamoxifen treatment.

## Methods

Adult male and female mice (defined as over 2 months of age) on a mixed C57Bl/6J and 129/SvEv background were housed in individually ventilated cages and fed with a regular chow diet. All animal work was carried out with a licence provided by the UK Home Office after local ethical and veterinary review. All mice described in this report were originally intended to provide controls for gene deletion experiments, or used to study the effects of gene deletion with Cre or CreER^13^. Mice given medetomidine anaesthesia and enrolled in experiments scheduled to persist for at least 7 days post-anaesthesia were included; in total, 100 animals were studied. Mice were injected intraperitoneally on two occasions with 0.5 mg tamoxifen (Sigma-Aldrich) dissolved in 85 µl or 250 µl vegetable oil (Sigma-Aldrich); this tamoxifen dose was chosen as a reported effective dose to induce genetic deletion in adult mice with *Cagg*-CreER^TM 13^. In some studies, mice were injected 7 and 3 days or 7 and 0 days before anaesthesia (together, 12 males and 17 females). In other studies, mice were injected 28 and 14 days prior to anaesthesia in an attempt to mitigate potential tamoxifen toxicity (32 males and 15 females). In both types of studies, tamoxifen-treated mice were compared with uninjected mice (18 males, 6 females). As per local recommended anaesthetic practice to induce deep anaesthesia prior to surgery, mice were then given 75 mg/kg of ketamine (Narketan; Vetoquinol) and 0.5 mg/kg of medetomidine (Dormitor; Vetoquinol), both diluted in sterile water. Intraperitoneal injection of 1 mg/kg atipamezole (Antisedan; Zoetis) was used to resolve medetomidine action. Mice were kept on a warming pad until fully recovered from sedation. Mice were returned to their home cages and monitored daily for 7 days after anaesthesia. Mice were monitored daily for clinical signs such as hunching and grimacing, mobility and signs of a hard or distended bladder. Mice were euthanised by an appropriate schedule 1 method at the end of the experiment or upon reaching their humane endpoints. Humane endpoints were defined as an inability to urinate, severely reduced mobility or signs of distress or pain as evidenced by the grimace scale. All data analysed here were obtained from mice used for pilot studies designed to test the effect of gene deletion on cardiovascular outcomes, including appropriate controls. Accordingly, the mice analysed here were pooled from several such pilot studies and did not themselves comprise a designed toxicity study; specifically, mice had not been randomised into uninjected and tamoxifen-treatment groups. All authors were aware of which groups mice had received tamoxifen treatment and which had not, both during weight measurements and recording of the onset of adverse effects, because the mice had originally been allocated to pilot studies testing the effect of gene deletion, rather than testing whether there is an interaction of tamoxifen treatment with subsequent anaesthesia. Statistical analysis was performed using GraphPad Prism v9.2. Survival curves were analysed for difference using a Log-rank (Mantel-Cox) test. P-values of ≤0.05% were considered significant.

## Results

Adult male and female mice were injected intraperitoneally with 0.5 mg tamoxifen on two occasions prior to anaesthesia with 75 mg/kg ketamine and 0.5 mg/kg medetomidine and the resolving anaesthesia using atipamezole (**Fig. 1A**). We unexpectedly observed that male mice injected with the second tamoxifen dose 0-3 days prior to induction of anaesthesia had a 50% incidence of adverse effects compared to 9.4% in mice that had received their second tamoxifen dose 14 days prior to anaesthesia, and male mice not given tamoxifen had no adverse effects (**Fig. 1B, C**). Severe adverse effects were seen in male mice expressing CreER and in CreER-negative control male mice.

**Figure 1:**
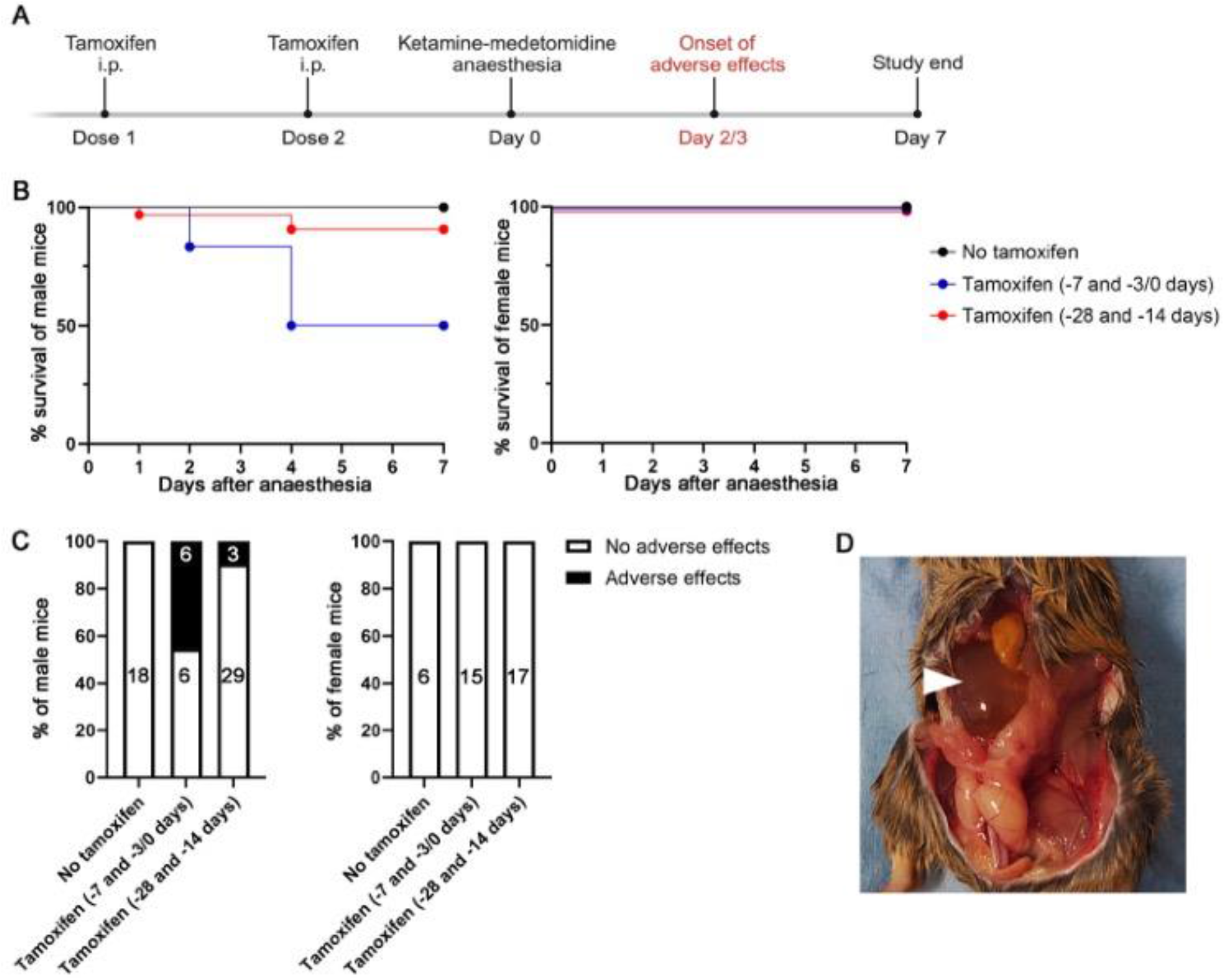
Tamoxifen injection followed by medetomidine-ketamine anaesthesia causes morbidity and mortality in male but not female adult mice. Prior to anaesthesia with ketamine-medetomidine, mice were either not injected (n = 18 males, n = 6 females) or injected intraperitoneally (i.p.) with 2 doses of tamoxifen (n = 44 males, n = 32 females). **(A)** Treatment and analysis timeline: Day 0 is defined as the day of anaesthesia with ketamine-medetomidine. In some studies, mice were injected with tamoxifen on day 7 (dose 1) and either on day 3 or 0 (dose 2) prior to anaesthesia (day -7 and -3/0). When the second tamoxifen dose was given on day 0, tamoxifen was given prior to anaesthesia. In other studies, mice were injected intraperitoneally with tamoxifen on days 28 (dose 1) and 14 (dose 2) prior to anaesthesia (day -28 and -14). All studies were terminated 7 days after anaesthesia. **(B)** Survival curves of male and female mice up to 7 days after anaesthesia. Male mice injected with tamoxifen on day -7 and -3/0 prior to anaesthesia had significantly reduced survival (6 out 12 male mice lost) compared to uninjected (no tamoxifen) male mice (0 out of 18 male mice lost); P = 0.0009. Male mice injected with tamoxifen on day -28 and -14 prior to anaesthesia had a lower rate of adverse effects (3 out of 32 male mice lost); this observation was not statistically significant at the number of mice included in our studies (P = 0.1861). No female mice were lost. **(C)** Percentage of male and female mice with adverse versus no adverse effects within 7 days of anaesthesia, after mice had received no tamoxifen or tamoxifen on day -7 and -3/0 or day -28 and -14 schedule. Black numbers on the white bars indicate the number of mice without adverse effects and white numbers on the black bars indicate the number of mice with adverse effects. **(D)** Example of a male mouse culled due to severe adverse effects, sprayed with ethanol before autopsy to prevent fur shedding; the arrowhead indicates a distended bladder.

Moreover, female mice showed no obvious adverse effects, regardless of their tamoxifen dosing schedule (**Fig. 1B, C**). Mice found at their humane endpoints presented with an inability to urinate when scruffed. Of the male mice euthanised at their humane endpoint, 3 mice were analysed post-mortem for gross abnormalities, and all were found to have distended bladders (**Fig. 1D**). Some mice were found dead, thereby exceeding the pre-specified humane endpoint of the study, requiring the filing of a report sent to the named animal welfare office, local veterinarian and legislator. Specifically, we observed a significantly decreased survival rate up to 72 hours post-anaesthesia in male mice injected with tamoxifen compared to male mice not injected with tamoxifen (P>0.001) (**Fig. 1B**). By contrast, female mice did not have decreased survival after tamoxifen dosing followed by anaesthesia (**Fig. 1B**).

## Discussion

Our study has established that the combination of tamoxifen injection with the anaesthetic agent medetomidine increased mortality in male mice 24-72 hours post-anaesthesia. Medetomidine is comprised of a combination of 2 enantiomers, dexmedetomidine and levomedetomidine, whereby dexmedetomidine is thought to be the active form and typically used at half the dose of medetomidine ^10^. Both medetomidine and dexmedetomidine have been suggested to be comparable in anaesthetic effect, and each needs to be combined with ketamine for deep anaesthesia ^10^. Even though medetomidine and dexmedetomidine are commonly used in both veterinary medicine and research with live mice ^10^, a literature search after observing adverse effects identified two published studies reporting that medetomidine and dexmedetomidine cause mortality in C57Bl/6J male mice at an incidence of 3% and 67%, respectively ^11, 12^. In agreement with the lower incidence of 3% mortality with medetomidine compared to dexmedetomidine, we did not observe any deaths in the small cohort of male mice that had received ketamine-medetomidine but no other intervention. By contrast, we observed increased mortality in 50% of male mice treated with tamoxifen within 3 days and in 9.4% of male mice treated with tamoxifen within 14 days of receiving medetomidine. These findings suggest that tamoxifen exacerbates medetomidine toxicity and that a shorter interval between administering both drugs is the most detrimental.

Medetomidine has been used in a range of domestic animal species for veterinary semen collection with a catheter under anaesthesia and has been associated with increased sperm counts ^14, 15^. In agreement, the prior study reporting increased mortality in male mice linked this to the formation of seminal coagulum that blocked the urethra ^12^, presumably because ejaculation was stimulated but the ejaculate was not released or manually removed, as is the case in veterinary practice for semen collection. As previously described for medetomidine-anaesthetised male mice in a study not using tamoxifen ^12^, the medetomidine-anaesthetised male mice treated with tamoxifen in our study also had distended bladders on post-mortem. Nevertheless, it is unclear why tamoxifen treatment increased the incidence of this medetomidine-induced adverse effect. As tamoxifen treatment in male mice decreases the proportion of mature elongated sperm cells at the expense of rounded, immature spermatogonia in the Sertoli cells ^16^, it is conceivable that immature sperm cells released into the seminal fluid may increase the incidence of urethral plug formation.

It cannot formally be excluded that elements of the anaesthetic regimen other than medetomidine increased mortality in tamoxifen-treated mice, or that mechanisms other than seminal coagulum are responsible; however, both possibilities seem unlikely, because female mice did not show mortality in our study. Moreover, a prior report showed that medetomidine toxicity in male mice was mitigated by replacing medetomidine with xylazine, whilst retaining ketamine and atipamezole in the anaesthetic protocol ^12^. Further re-analysis of existing or ongoing studies carried out in other labs with other anaesthetic regimens would be required to determine whether tamoxifen can also exacerbate the side effects of other anaesthetic regimens. We trust that the report would precipitate such analysis to improve animal welfare in Cre-LoxP experiments.

Notably, the mortality and morbidity associated with tamoxifen use prior to medetomidine-based anaesthesia decreased with a longer duration between the last tamoxifen dose and the induction of anaesthesia. Thus, mortality in male mice was reduced from 50% to 9.4% when the last dose of tamoxifen was administered 14 days before anaesthesia, as opposed to 0 or 3 days before anaesthesia. Nevertheless, we consider a 10% mortality incidence unacceptable. An even longer duration between tamoxifen administration and induction of anaesthesia may further reduce the mortality rate but may be unsuitable for some experimental models due to recovery from the intended purpose of tamoxifen administration, which is to induce gene deletion. Thus, a population of recombination-resistant cells may have an advantage over recombined cells and out-compete them (e.g., Fantin et al. Blood, 2013^17^). Therefore, we suggest that alternative injectable anaesthetic formulations not containing medetomidine should be used when using male mice, particularly in combination with tamoxifen administration to induce gene deletion, and when inhalation anaesthesia is unsuitable.

## Animal welfare implications

This report seeks to document and alert other researchers to the adverse effect of tamoxifen administration prior to medetomidine anaesthesia in male mice. The significant morbidity and mortality we have observed constitute ethical reasons prevent us including additional animals in a thoroughly designed experimental study with larger group sizes. Nevertheless, our report should significantly impact animal welfare by prompting researchers to avoid medetomidine and dexmedetomidine for anaesthesia of tamoxifen-treated mice in genetic recombination studies using Cre-LoxP. This is particularly important, because both tamoxifen and medetomidine are commonly injected into mice for *in vivo* experiments to study gene function in injury and disease models seeking to provide novel information relevant to human health, yet, alternative anaesthesia protocols are available for which adverse effects have not been reported.

## Data availability

Study data are stored securely at UCL and are available on request from C.R.

## Acknowledgements

V.S.R., L.D. and C.R. were supported by research grants from the British Heart Foundation [PG/19/37/3439, FS/18/65/34186] and Wellcome [205099/Z/16/Z]. The funders had no role in the design, analysis, and reporting of the study.

## References

1. Nagy A. Cre recombinase: the universal reagent for genome tailoring. Genesis 2000; 26: 99–109.

2. Feil R, Brocard J, Mascrez B, et al. Ligand-activated site-specific recombination in mice. Proceedings of the National Academy of Sciences of the United States of America 1996; 93: 10887–10890. DOI: 10.1073/pnas.93.20.10887.

3. Metzger D, Clifford J, Chiba H, et al. Conditional site-specific recombination in mammalian cells using a ligand-dependent chimeric Cre recombinase. Proceedings of the National Academy of Sciences of the United States of America 1995; 92: 6991–6995. DOI: 10.1073/pnas.92.15.6991.

4. Loonstra A, Vooijs M, Beverloo HB, et al. Growth inhibition and DNA damage induced by Cre recombinase in mammalian cells. Proceedings of the National Academy of Sciences of the United States of America 2001; 98: 9209-9214. 2001/08/02. DOI: 10.1073/pnas.161269798.

5. Bersell K, Choudhury S, Mollova M, et al. Moderate and high amounts of tamoxifen in alphaMHC-MerCreMer mice induce a DNA damage response, leading to heart failure and death. Dis Model Mech 2013; 6: 1459-1469. 2013/08/10. DOI: 10.1242/dmm.010447.

6. Brash JT, Bolton RL, Rashbrook VS, et al. Tamoxifen-Activated CreERT Impairs Retinal Angiogenesis Independently of Gene Deletion. Circ Res 2020; 127: 849-850. 2020/07/09. DOI: 10.1161/CIRCRESAHA.120.317025.

7. Rashbrook VS, Brash JT and Ruhrberg C. Cre toxicity in mouse models of cardiovascular physiology and disease. Nat Cardiovasc Res 2022; 1: 806-816. 9/9/2022. DOI: 10.1038/s44161-022-00125-6.

8. Sun MR, Steward AC, Sweet EA, et al. Developmental malformations resulting from high-dose maternal tamoxifen exposure in the mouse. PLoS One 2021; 16: e0256299. 2021/08/18. DOI: 10.1371/journal.pone.0256299.

9. Ved N, Curran A, Ashcroft FM, et al. Tamoxifen administration in pregnant mice can be deleterious to both mother and embryo. Laboratory animals 2019; 53: 630–633. DOI: 10.1177/0023677219856918.

10. Burnside WM, Flecknell PA, Cameron AI, et al. A comparison of medetomidine and its active enantiomer dexmedetomidine when administered with ketamine in mice. BMC Vet Res 2013; 9: 48. 2013/03/19. DOI: 10.1186/1746-6148-9-48.

11. Cagle LA, Franzi LM, Epstein SE, et al. Injectable Anesthesia for Mice: Combined Effects of Dexmedetomidine, Tiletamine-Zolazepam, and Butorphanol. Anesthesiol Res Pract 2017; 2017: 9161040. 2017/02/18. DOI: 10.1155/2017/9161040.

12. Wells S, Trower C, Hough TA, et al. Urethral obstruction by seminal coagulum is associated with medetomidine-ketamine anesthesia in male mice on C57BL/6J and mixed genetic backgrounds. J Am Assoc Lab Anim Sci 2009; 48: 296-299. 2009/05/30.

13. Brash JT, Denti L, Ruhrberg C, et al. VEGF188 promotes corneal reinnervation after injury. JCI Insight 2019; 4 2019/11/02. DOI: 10.1172/jci.insight.130979.

14. Madrigal-Valverde M, Bittencourt RF, Ribeiro Filho AD, et al. Quality of domestic cat semen collected by urethral catheterization after the use of different alpha 2-adrenergic agonists. J Feline Med Surg 2021; 23: 745-750. 2020/11/19. DOI: 10.1177/1098612X20973183.

15. Zambelli D, Cunto M, Prati F, et al. Effects of ketamine or medetomidine administration on quality of electroejaculated sperm and on sperm flow in the domestic cat. Theriogenology 2007; 68: 796-803. 2007/07/31. DOI: 10.1016/j.theriogenology.2007.06.008.

16. Patel SH, O’Hara L, Atanassova N, et al. Low-dose tamoxifen treatment in juvenile males has long-term adverse effects on the reproductive system: implications for inducible transgenics. Sci Rep 2017; 7: 8991. 20170821. DOI: 10.1038/s41598-017-09016-4.

17. Fantin A, Vieira JM, Plein A, et al. NRP1 acts cell autonomously in endothelium to promote tip cell function during sprouting angiogenesis. Blood 2013; 121: 2352–2362. DOI: 10.1182/blood-2012-05-424713.

